# Human genetic disease is greatly influenced by the underlying fragility of evolutionarily ancient genes

**DOI:** 10.1101/558916

**Authors:** Alexandra Martin-Geary, Mark Reardon, Benjamin Keith, May Tassabehji, David L. Robertson

**Affiliations:** Division of Evolution and Genomic science, University of Manchester, Oxford Rd, Manchester M13 9PL; MRC-University of Glasgow Centre for Virus Research, Glasgow G61 1QH

## Abstract

The ability to predict disease association in human genes is enhanced by an evolutionary understanding. Importantly genes linked with heritable disease, particularly dominant disorders, tend to have undergone duplication in our early vertebrate ancestors, with a strong asymmetric relationship between disease-association within duplicate/paralog pairs. Using a novel phylogenetic approach, alongside a whole-genome comparative analysis, we show that contrary to the accepted compensatory model of disease evolution, the majority of disease-associations reside with the more evolutionary constrained gene, inferred to most closely resemble the progenitor. This indicates that the strong association between paralogs, specifically ohnologs, and dominant disorders is often a consequence of a mechanism through which pre-existing dosage sensitive/haploinsufficient genes are successfully duplicated and retained. Heritable disease is thus as much a consequence of the fragility of evolutionarily more ancient genes as compensatory mechanisms. From these findings, we demonstrate the utility of a new model with which to predict disease associated genes in the human genome.

## Introduction

More than 150 years since Gregor Mendel’s findings were first published, the role of evolution and heritability in human genetics, and specifically disease association, is still being elucidated. We now know that whilst Mendel’s foundation of dominant versus recessive is broadly correct, a myriad of factors can contribute to both penetrance and severity of the resulting phenotype, but also that complex genetic interactions, where the interplay between multiple genes results in a trait, rather than the simple ‘one gene-one trait’ model can and does occur (Cooper et al., 2013). In terms of mono-genic disease association, a striking association has been found to exist between genes duplicated in vertebrate evolution and disease with 80% of human heritable disease genes residing in a paralog in the human genome (Dickerson and Robertson, 2012). Why does this strong one gene to phenotype relationship exist with genes that have a duplication event in their history?

While ‘duplicability’ of a gene can be influenced by genomic context, such as sequence composition and chromosomal location, leading to much of the copy number variation observed within the human genome (Schuster-Böckler et al., 2010; Truty et al., 2018), differential retention biases have played a significant role in the landscape of paralogs observed in genomes (Hakes et al., 2007). Notwithstanding stochastic sampling processes in evolution associated with smaller population sizes (Lynch and Conery, 2003), this will often be a result of factors associated with gene product fitness. Whole genome duplication in particular, is hypothesized to increase the chance of retention of dosage-threshold sensitive genes, as a consequence of maintenance of stoichiometric balance (Papp et al., 2003), which, due to negative impact on the system, would be very unlikely to be retained within the context of small-scale duplication (SSD) events (Makino and McLysaght, 2010). Furthermore, it has been proposed that whole-genome duplication (WGD) events confer an immediate fitness benefit to the organism by reducing expression ‘noise’ (Pires and Conant, 2016). While copy-number associated duplicates tend to be refractory for the retention of dosage-sensitive paralogs (Makino and McLysaght, 2010).

These studies demonstrate the imperative of having a mechanistic and evolutionary understanding of the processes by which genes arise and are retained, in order to disentangle the relationship between gene duplication and disease (Innan and Kondrashov, 2010; Opazo and Zavala, 2018). Exemplifying this perspective is the strikingly strong association observed between genes which have undergone duplication in vertebrate evolutionary history, particularly WGDs and heritable human disease (Makino and McLysaght, 2010). Following a ‘compensation model’, where duplicates contribute to redundancy (Gu et al., 2003), we hypothesised that this relationship was due to the accumulation of otherwise deleterious mutations in the context of duplication events (Dickerson and Robertson, 2012), with disease associations emerging as new genes arise and are retained, i.e., duplication introducing disease potential by ‘masking’ of otherwise deleterious mutations. Singh *et al* (Singh et al., 2012) presented a modified version of the compensation model, in which the compensation for deleterious variation, in particular for dominant disorders, is ‘locked in’ by the WGD event, and subsequently neither gene in the duplicate pair can be lost without severe consequences to fitness.

What is often neglected, however, is the asymmetric relationship of disease to paralog pairs (Dickerson and Robertson, 2012), which must be considered in any analysis. Due to differential pressures, following a duplication event it is rarely the case that the two paralogous genes will remain identical indefinitely. Over time the accumulation of variants in either or both of the duplicates leads to divergence. This divergence can result in differing functions of the two genes, sub- and neo-functionalization, or pseaudoginization amongst others. It can also lead to disease. The relative proportions of these outcomes is a contentious subject, with conflicting theories surrounding not only differential retention, but also the likelihood of and degree to which evolutionary asymmetry may occur (Pachter, Lior, 2015). Fundamentally, the discussion revolves around two arguments: the first, that paralog pairs tend to show asymmetry where the less constrained copy is likely to be harmless to the organism, presented by Ohno *et al* (Ohno, Susumu, 1970) and the second, proposed by Force *et al* (Force et al., 1999) that evolution following duplication is unlikely to be asymmetric. The current consensus agrees with Ohno’s proposition that evolutionary asymmetry does exist (Kellis et al., 2004), although the statistical measurement employed remains contested (Pachter, Lior, 2015), and may be dependent on the degree of dosage sensitivity within any given paralog pair (Tasdighian et al., 2017).

In contrast to this it has been hypothesized that, following duplication, redundant genes confer robustness through the provision of a functional paralog, which acts as a ‘buffer’ to otherwise deleterious disruptions to its partner gene (Gu et al., 2003; Hakes et al., 2007; Hsiao and Vitkup, 2008; López-Bigas and Ouzounis, 2004; Rice and McLysaght, 2017). It is proposed that this dilution of deleteriousness permits the retention of disease-associated genes, which would otherwise be subject to purifying selection. Given this ‘compensation’ model, we would predict that disease-associated mutations would mostly be found on the less constrained, longer branch within the gene pair as the relaxation of purifying selection will permit accumulation of slightly deleterious mutations, i.e., non-lethal disease-causing mutations (Dickerson and Robertson, 2012). Thus, explaining the connection between paralogs and heritable disease.

Another important variable is the haploinsufficient nature of genes and its link to dosage and disease (Veitia and Birchler, 2010), i.e., those where a single functioning copy of the diploid gene alone is unable to support the wild-type function. Haploinsufficiency is strongly associated with gene duplication (Kondrashov and Koonin, 2004; Papp et al., 2003; Pires and Conant, 2016), and has generally been found to have a greater number of paralogs than haplosufficient genes. This is likely due to the need to retain genes of this kind following duplication events/dosage effects as they have a greater likelihood of negatively impacting on the system, leading to stoichiometric imbalance should differential copy numbers arise (Makino and McLysaght, 2010).

To investigate the association between evolution, the diploid nature of human genes and human disease further, we parsed these sets of trees using a novel phylogenetic method for gene dating and found that heritable disease genes tend to be evolutionary ancient and associated with gene duplication, and in the case of dominant disorders this is due to WGD being the mechanism for duplicating haploinsufficient genes. Confirming this we find disease-associated mutations tended to fall on the shorter constrained branch in paralog pairs, and are thus more likely to be associated with the ancestral function. We discuss these findings and the implication that disease states are due to a pre-existing fragility within the ancestral genome, rather than the product of later duplication events.

## Results

Using a novel phylogenetic method of dating genes (SI), we were able to assign singleton and duplicate gene ages and compare properties of disease and non-disease genes. From this analysis, we have been able to determine that both dominant and recessive disease-associated genes, as identified within the OMIM database, tend to be relatively evolutionarily ancient (Figure 1A). Significantly, the majority of these genes arose either with the two rounds of whole genome duplication (WGD, ~441 MYA in our Euteleostomi/fish ancestors before the split between cartilaginous and bony vertebrates) or predate them, with relatively few disease-associated genes arising thereafter. The cumulative frequency of heritable disease-associated genes within the human genome (Figure 1B) reveals a relative plateau in the introduction of obviously disease-associated genes in more recent evolutionary history, the onset of which coincides with the most recent round of WGD ~441 million years ago (MYA). This is consistent with the proposal that there has been a relatively low rate of disease associated gene introduction following the last round of WGD, with the majority of disease being evolutionarily ancient.

**Fig. 1.**
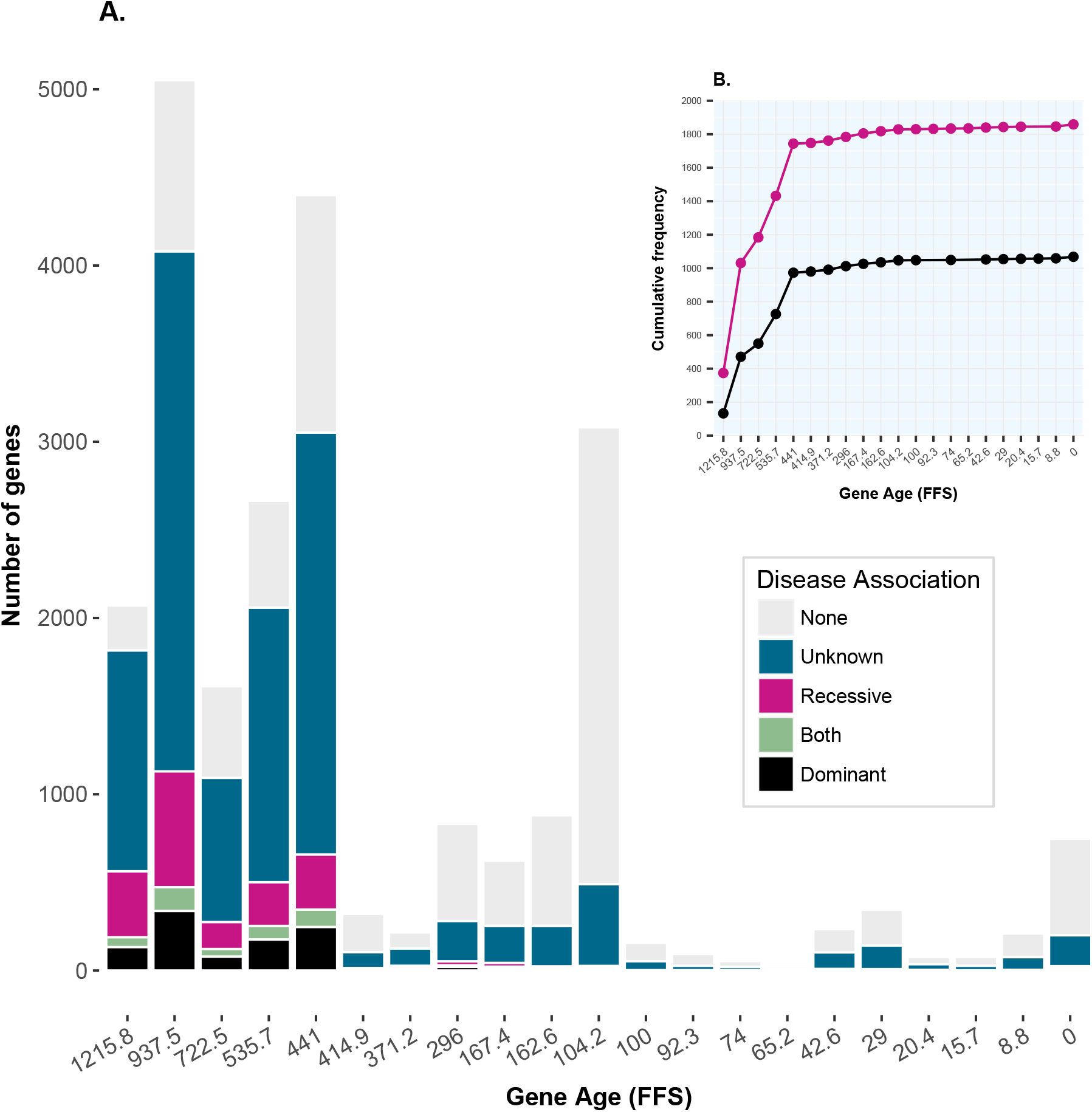
Bar chart showing numbers of genes with differing disease states during evolutionary past (inferred from taxonomic levels), showing a clear trend in older genes being disease-associated, and a spike at 104.2 MYA in our Eutheria/mammalian ancestors, likely representing the branching of placental mammals which occurred at this time point (A). Inset: cumulative frequency of disease genes over time for dominant and recessive disease associated genes (B).

Support for this finding of a high association between ancient genes and fragility is provided by the results of our analysis of gene age and haploinsufficiency (HI), using the Decipher haploinsufficiency scores (Firth et al., 2009) (Figure 2). Genes falling into the ‘true’ haploinsufficient decile, as defined by Decipher (Firth et al., 2009), are almost exclusively genes arising at, or prior to, the most recent round of WGD (441 MYA), with only the only outliers arising from the time-point immediately after WGD. While the seminal paper Makino and McLysaght (2010) also looked at the link between dosage and ohnologs, they did not directly measure this using haploinsufficiency rather assuming HI to be a property of the ohnologs.

**Fig. 2.**
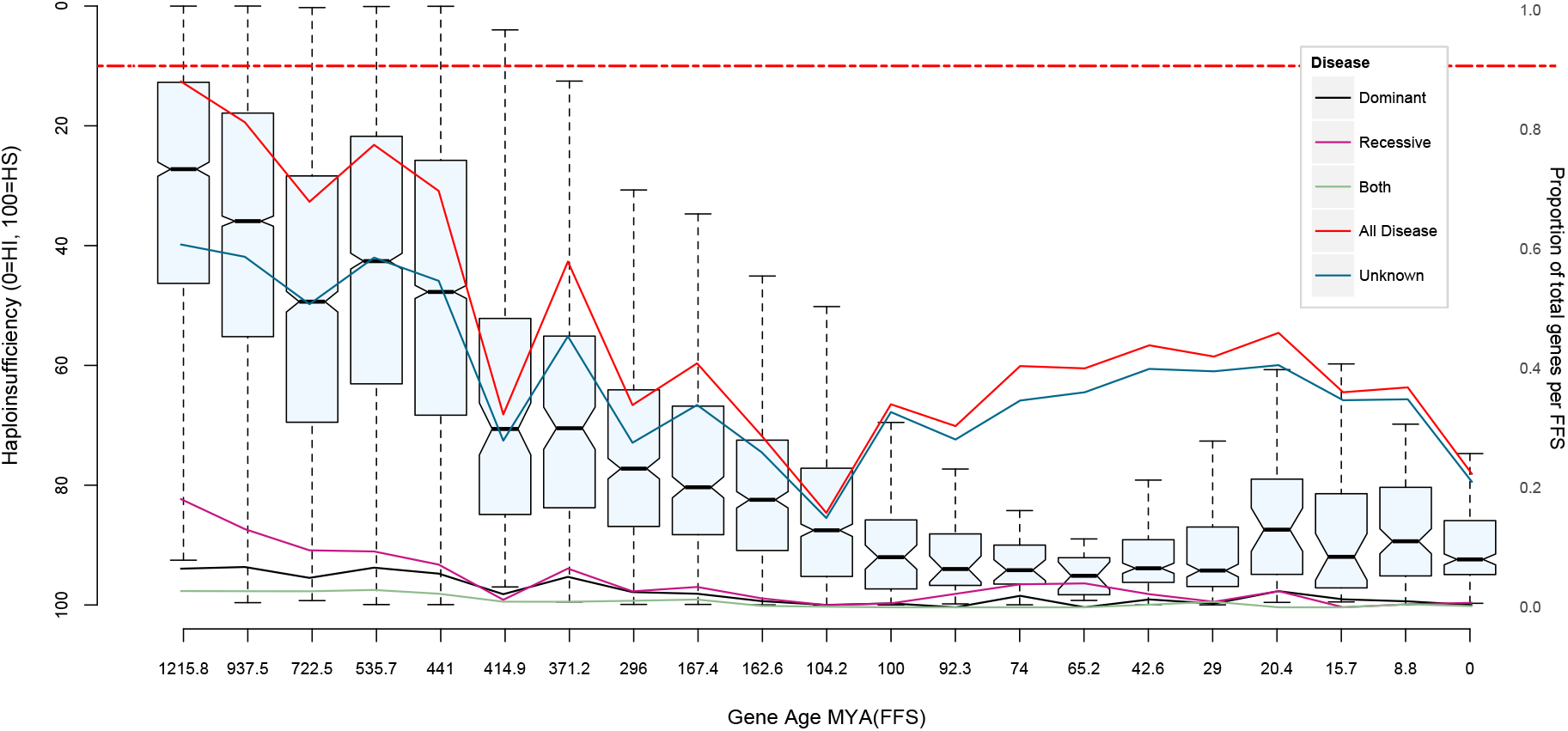
Notched box and whisker plot showing haploinsufficiency of genes within each gene age bin, where 0 is the most, and 100 is the least haploinsufficient (HI). The dashed red line shows the conservative cut-off of haploinsufficiency proposed by Decipher (Firth et al., 2009), above which they predict ‘true’ haploinsufficient genes to reside. Overlaid is a line graph plotting the normalized frequency of disease genes in each age, between 0 and 1, arising at each time point.

To test the significance of this association we investigated the relationship between disease, haploinsufficiency, paralog status and gene age (Figure 3). The multiple correspondence analysis (MCA) of these four features demonstrates a strong relationship between disease, paralog status, gene age, and haploinsufficiency, in particular, for duplicated genes with dominant disease-associations. This indicates haploinsufficiency is providing the underlying structure of their interactions.

**Fig. 3.**
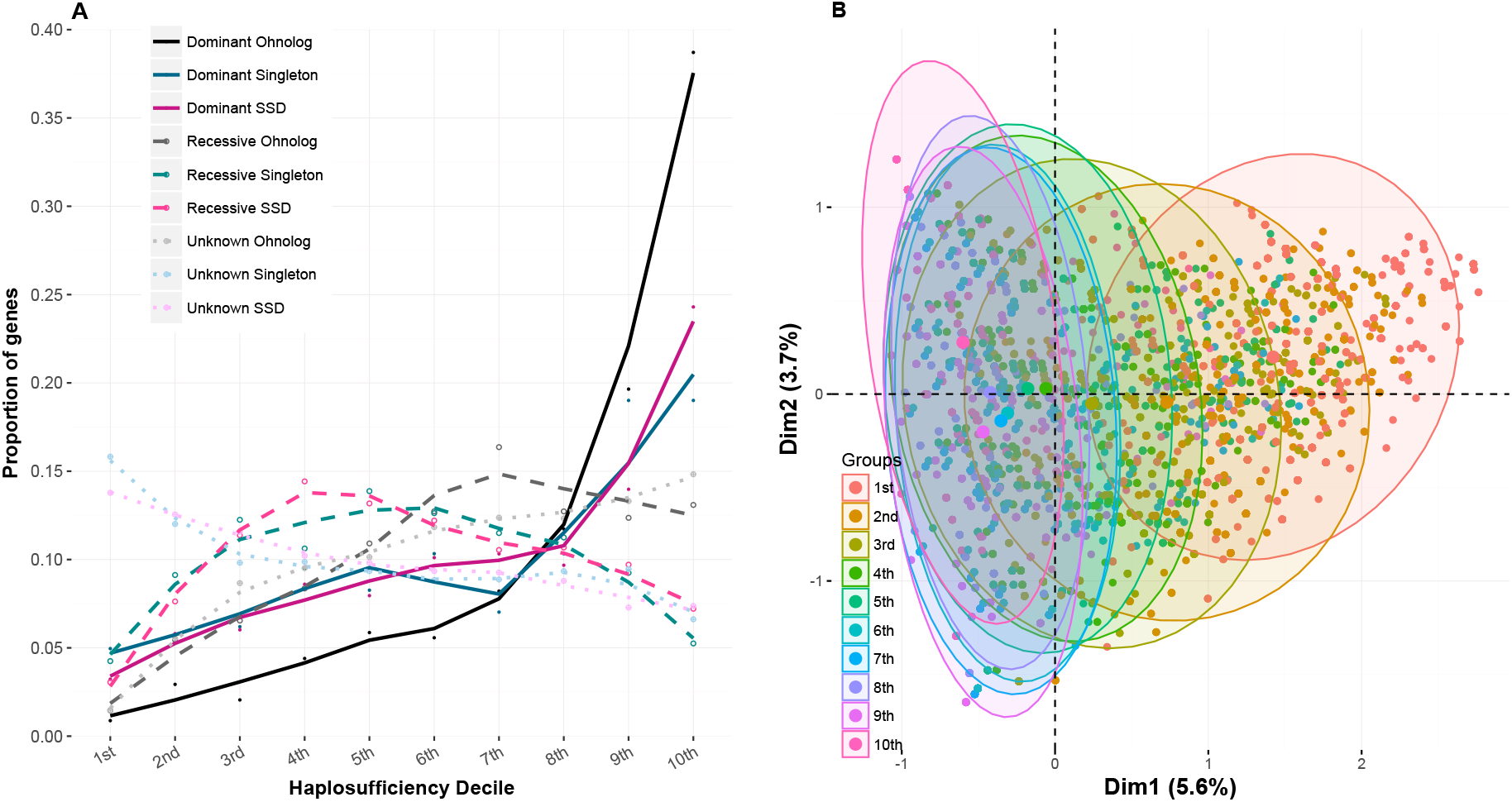
Total proportion of genes from various disease and duplication states which reside in each haploinsufficiency decile (A); 1 being the least and 10 being the most haploinsufficient; solid lines show genes with dominant disease-associations, and dashed lines show those with recessive disease-associations. Multiple correspondence analysis of haploinsufficiency, gene age, paralog status and disease status (B); the 10 haploinsufficiency deciles are highlighted by the coloured ellipses.

We tested the hypothesis that disease-association preferentially tracks to the more diverged gene within any paralog pair. In order to do this, we parsed sets of gene trees to establish relative branch lengths of genes in both ohnolog and SSD pairs. This novel method provides a way of dating genes, accounting for their duplication histories by comparing relative branch lengths and ordering of Ensembl gene families (see supplementary method). This was further refined by establishing disparity of function by comparing associated GO terms between genes within pairs, and subsequently refining to genes pairs with observed functional asymmetry. Confirming our previous analysis (Dickerson and Robertson, 2012), the majority of disease associations fall on just one of the paralogs partners (78%/1510 pairs). while in only 433 cases (22%) are both paralogs disease associated. Contrary to expectations, however, more of the pairs containing just one gene associated with disease associate with the constrained, i.e., shorter branch (56%/840genes)), than the longer/evolutionary less constrained branch (44%/710 genes) (Figure 4A) as predicted by the disease being due to disease mutations accumulating as a consequence of relaxed selection (Dickerson and Robertson, 2012), confirmed to be statistically significant (Chi-squared tests, P-value =2.2E-16). This indicates that disease is often a consequence of mutations disrupting the ancestral function, i.e., inferred to be retained by the more conserved paralog.

**Fig. 4.**
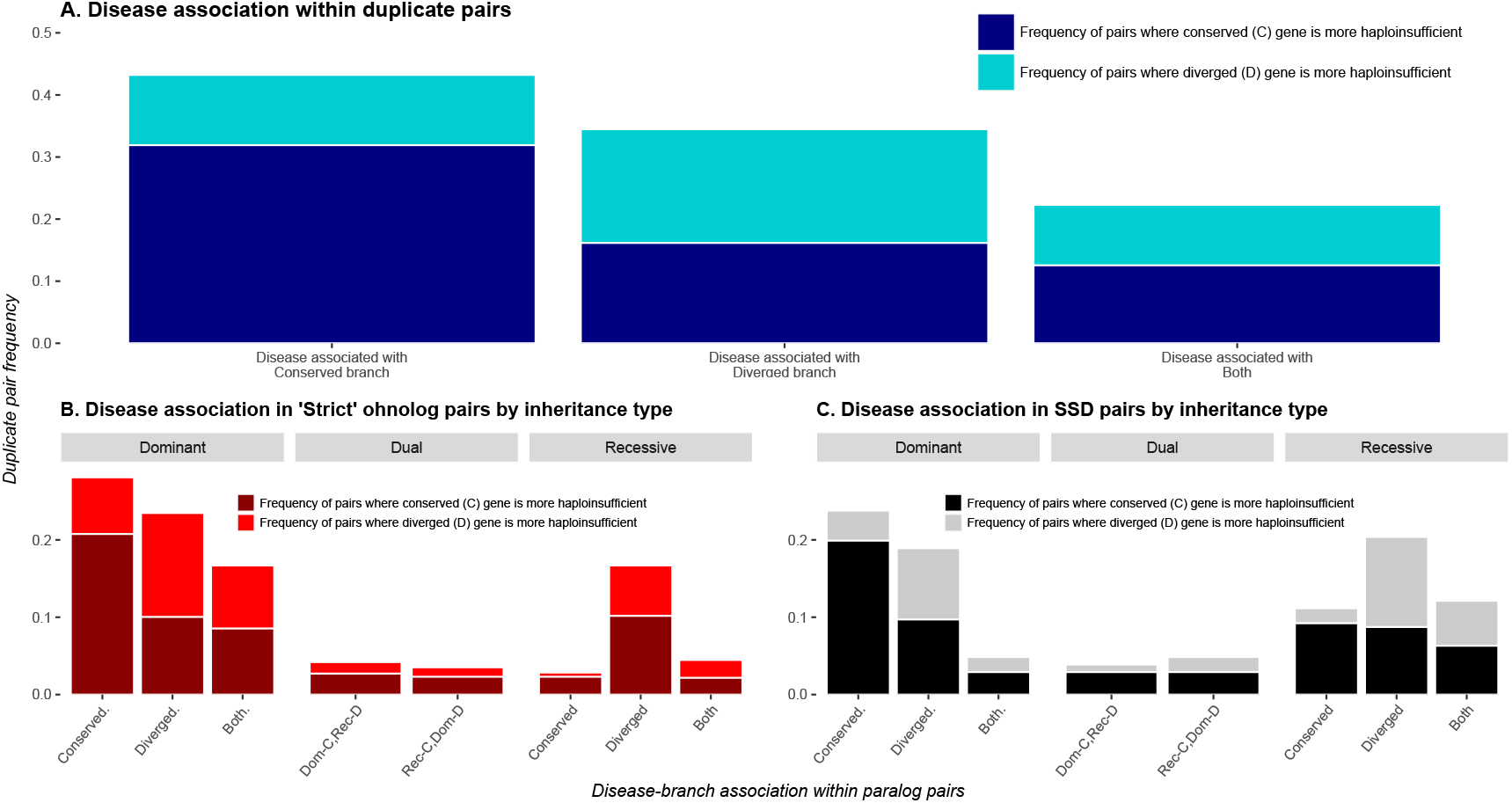
Bar charts showing asymmetry between paralog pairs with differing disease-associations, evolutionary divergence, and haploinsufficiency for all duplicate pairs (A), ohnolog pairs including ohnologs with a later duplication by inheritance type (B), and small-scale duplication (SSD) pairs by inheritance type (C). Dark bars correspond to pairs where the most conserved gene is also the most haploinsufficient; light bars are pairs where the more diverged gene is the more haploinsufficient. Disease-associations are shown on the x-axis.

For the diseases associated with the conserved paralog these are over-represented for the more haploinsufficient genes (74%), further confirming the role of haploinsufficient genes in disease. This focus on asymmetry shows that disease propensity is most frequently associated with a combination of both haploinsufficiency and conservation, wherein the disease associated paralog retains a greater sequence similarity to the ancestral gene than its partner.

When adding heritable disease type to this analysis, disease, particularly dominant disease, being associated with the conserved branch was again the case, with more tending to be haploinsufficient (Figures 4B & C). The major role of haploinsufficiency is supported by gene pairs that possess a dominant disease-association only in the more diverged gene (Dominant D, Figure 4), whilst these occur considerably less frequently than pairs whose disease association resides in the more conserved gene, when they do, it is predominantly the case that the disease gene also ranks more highly in the haploinsufficiency scale. Indeed there is a highly significant trend in the increase in the number of pairs where disease is present in only one gene, which is also both more conserved and has a greater haploinsufficiency score (SI table S2). It must also be noted that, whilst the profiles are broadly similar in both ohnologs (Figure 4B), and SSDs (Figure 4C) the exception is in the relative proportions of recessive disease, which highlights the propensity of ohnologs to be more frequently associated with dominant disorders.

These results indicate it is frequently the paralog that retains the ancestral function that has a tendency towards fragility and therefore disease. If this is indeed the case, although not over-represented for any disease-association (Singh et al., 2015), we would still expect to see the same pattern for SSDs, and this is what we find (Figure 4C). As with WGDs, for SSDs associated with dominant disorders, it is again the constrained/shorter branch that tends to have a higher association with haploinsufficiency and, unlike WGDs, there is also a high association with recessive disorders. This asymmetric relationship confirms that disease tends to be associated with the more constrained/older gene function.

## Discussion

Our findings demonstrate that the association between ohnologs and dominant disease exists for the most part due to their tendency to be haploinsufficient, a feature most commonly associated with genes arising prior to the two known WGD events in our early vertebrate ancestors (Figure 2). The overrepresentation of disease genes within this set then arises because WGD is the main process by which haploinsufficient genes can be both duplicated and have a high probability of being retained. This provides an understanding of why so many dominant diseases exist within human populations.

This is consistent with a recent study by Diss *et al* (Diss et al., 2017) who investigated the impact on the system caused by the introduction of interdependent paralogs and show differential dependence in the model organism yeast, suggesting that duplication has introduced fragility and not increased robustness as expected. This supports the hypothesis of equivalent divergence (and therefore absence of asymmetry) presented by Force *et al* (Force et al., 1999), as disruption of interdependent subfunctions of one of the duplicated paralogs in a subfunctionalised pair would produce a deleterious phenotype.

The evolution of complexity has been facilitated by gene duplication mechanisms, particularly WGD events, which have facilitated the introduction of repurposable genetic material, despite the pre-existing fragility of more evolutionarily ancient functions. Ohnologs are a source of functional divergence (Acharya and Ghosh, 2016) as they increase the probability of a dosage-dependent gene being retained, the later elongation of branch length is indicative of novel function which, in terms of disease, is likely either to be associated with complex disease or no disease. That only one of the duplicates retains dosage-sensitivity is in line with theoretical work that incorporates retention bias due to dosage constraints (Teufel et al., 2016).

There are, however, a small proportion of genes which, whilst likely to fit our criteria of greater constraint and haploinsufficiency will never be observed to be disease associated. In the case of these genes this is due to the fundamental inviability which would be introduced as a result of their disruption, and which, as highlighted by previously observed links between essentiality, developmental processes and ohnologs (Makino et al., 2009), are likely to be overrepresented in genes which have resulted from a WGD event.

The significance of our finding that fragile, heritable-disease associated genes are evolutionary more ancient can assist in a predictive capacity. This is highlighted when applied to a newly identified disease set obtained from a recent study by Bastarache *et al* (Bastarache et al., 2018), who presented a new method using phenotype risk scores to identify novel Mendelian disease variants. They identified 18 variants in 16 genes leading to monogenic disorders. Using these genes as a test set, we predicted that they would predominantly be duplicates, and that those duplicates would have greater haploinsufficiency and shorter branch length when compared with their paralog partner. We found that of the 16 genes, 12 were duplicates: 9 ohnologs (from varying stringency criteria) and 3 SSDs (supplementary results table S2). Within this set, five of the newly-established disease genes had both the shortest branch length and highest haploinsufficiency scores, three had the shortest branch length and lowest haploinsufficiency scores, three had the longest branch length and highest haploinsufficiency scores, and only one gene had the longest branch and lowest haploinsufficiency score. This validates our finding that disease is most commonly associated with the more conserved gene in a pair. However, in cases where it is the divergent gene that is associated with disease, this is almost always the gene with a greater haploinsufficiency score.

By expanding our understanding of haploinsufficiency and paralog-associated disease asymmetry within duplicated gene pairs, we have demonstrated that the observed enrichment of dominant disease-association is an artifact of pre-existing haploinsufficiency and any subsequent disease states are due to this, rather than the somewhat counterintuitive hypothesis that it is dominant disease-association, so-called dangerousness, which leads to retention (Singh et al., 2012). It should be noted that the second round of WGD (Acharya and Ghosh, 2016) will add at least one additional disease-associated gene, so ohnologs do also add disease-prone genes to the genome. These haploinsufficient duplicates, which can persist due to the initial presence of a functionally identical partner, are then able to evolve away from their haploinsufficient disease-associated state. A high abundance of haploinsufficient disease-associated genes were already extant in the ancestral genome, and what we observe within ohnologs is an increase in genes with an on-going reduction in potential for monogenic disease associations compared with the pre-WGD genome.

In conclusion, the ancestral role of the progenitor genes was likely the provision of relatively more ‘core’ functions common to most life, which leads to the observed associations with haploinsufficiency, essentiality and, as a consequence, a tendency towards fragility, all of which are traits passed on to their duplicated ‘daughters’. The lack of obvious disease-association in more recent evolutionary history will be a consequence of a reduction in the importance of any one specific gene in the context of greater functional complexity.

## Methods

### Allocation of evolutionary ages to genes

To obtain the gene ages, the Ensembl IDs and gene types of all known genes were obtained from Ensembl (release 88, genome GRCh38) (Zerbino et al., 2018) with R (version 3.3.3) (R Core Team, 2016) using the biomaRt R package (version 2.28) (Durinck et al., 2009). The known gene IDs were used by a Perl script running on ActivePerl (version 5.20.2) that saved the approximate taxon age and relative phylogenetic branch length (based on protein sequence similarity) of all the speciation and duplication events for every known gene with a gene tree in Ensembl Compara, starting with each gene’s leaf node and recursively saving all the ancestor events in the gene’s tree up to the root event (SI Figure). The known gene IDs were grouped into paralog families using biomaRt and, together with the gene tree event ages and branch lengths, formed the input to a C# program (Microsoft Visual Studio Enterprise 2015, .NET version 4.6.2) that implements the FFS algorithm.

To obtain duplicate gene ages an algorithm, furthest from source, FFS, was devised. First the approach removes all speciation events apart from those at the start of gene histories and all duplication events marked as ‘dubious’, preserving the phylogenetic branch lengths from the removed events by adding them to the next youngest event. The gene tree events for each paralog family’s genes were then used to create a human-specific paralog tree, started by merging the oldest event (common to all genes in the family) from each gene’s event history and then continued by merging events where identical in age and branch length and branching otherwise. Non-bifurcating duplication events were then removed from the human-specific paralog trees, again preserving the removed events’ branch lengths by adding them to the next youngest event. This method of tree building recreates the human-specific duplication topology of the paralog family in Ensembl Compara, preserving branch lengths while removing events unnecessary for the allocation of duplication ages to genes.

The ages of the root node and internal duplication nodes in each paralog tree were then assigned to the gene leaf nodes according to the following algorithm, with node distances comprising the cumulative branch lengths between the node in question and the root node:

~~~
while there are genes in the paralog tree without ages…
      set the subject to be the undated gene furthest from the root
      for each duplication from the subject’s ancestor back up to the root…
       if the duplication has undated genes with shorter distance to root…
          subject takes the duplication’s age
       if subject is still undated…
          subject takes the root’s age
~~~

This algorithm has the effect of assigning ages from duplications to genes such that a gene’s FFS-assigned age is influenced by the degree of protein sequence similarity between the gene and the inferred ancestral gene at the root of the paralog tree. The root duplication/speciation node’s age is assigned to two genes in contrast to internal duplication nodes’ ages which are assigned to exactly one extant leaf gene node (triplication node ages would be assigned to two gene nodes, etc). In addition to paralog similarity to the inferred ancestral sequence, FFS takes account of all duplications in a paralog family. A small number of genes were found, wherein there had been no divergence since their most recent shared duplication event, these genes were defined as ambiguous and omitted from the current analysis.

The FFS approach also emits the IDs, names and gene-to-parent branch lengths of the ‘terminal paralog pairs’, defined as the gene pairs that have a duplication as their parent node. For each terminal paralog pair, the normalised branch length asymmetry was calculated as the difference in branch lengths of the pair divided by the maximum gene distance to root in each paralog family (so that the asymmetries could be compared across all paralog families).

### Disease status, paralog status, and haplosufficiency

In addition to gene ages, data was gathered pertaining to disease association, gene age and paralog status of all ‘known’ genes. Data analysis was performed using a combination of Perl 5, version 18, subversion 2 (v5.18.2) (Wall, 2000) and R version 3.3.2 (2016-10-31) (R Core Team, 2016).

Paralog status was calculated by cross-referencing output from the FFS approach with data pertaining to whole genome duplication status obtained from Singh *et al* (Singh et al., 2015) on 19/03/2018. It was decided when restricting to ‘ohnologs’ alone, to only include gene pairs which fulfilled the ‘strict’ criteria, having q-scores for both self-comparison, and outgroup of <0.01 (ibid), as these provide the clearest distinction between ohnolog and non-ohnolog genes in the human genome.

The FFS-dated genes were subsequently partitioned into paralog status based on their individual history of duplication, and presence in the ‘strict’, ‘intermediate’ or ‘relaxed’ synteny-based ohnolog lists. Those with an additional duplication event more recent than the second WGD event at 441mya were defined as ‘[ohnolog-type]/SSD’. Genes without any duplication events in their history were defined as ‘singleton’ and the remaining duplicated genes, not yet assigned a duplication type, were designated as ‘SSD’.

Data pertaining to disease association were obtained from OMIM in the form of the Genemap2 dataset (https://data.omim.org/downloads/dI1aeTBYTNet3PfqZqIS_w/genemap2.txt) obtained on 1/5/2018. Information contained therein used in the current analysis was compiled by OMIM from the following sources: Ensembl gene id – Ensembl; Phenotype data – OMIM. A text-based search was performed on this file to establish, firstly, if each gene in our ‘known’ set existed within the OMIM dataset, and if so, whether their association was with dominant, recessive or unknown disease inheritance, subsequent assignment of disease status was conducted according to these findings. For rare instances where genes had both dominant and recessive associations their status was defined as ‘both’ and genes in the ‘known’ list, not present in Genemap2, were defined as ‘none’.

Haploinsufficiency (HI) scores (Huang et al., 2010) were obtained from Decipher (Firth et al., 2009) (Haploinsufficiency Predictions Version 3 bed file) on 10/03/2018. Due to the fact that the entries in this file listed genes using HGNC identifiers, to bring them in line with the Ensembl naming convention used in the analysis it was necessary to cross reference gene names with their Ensembl counterparts. To this end Ensembl GRCh38.p7 gene names and HGNC IDs were obtained from Ensembl BioMart (Zerbino et al., 2018) on 03/05/2018, and, where possible HGNC IDs substituted in the Haploinsufficiency data, for Ensembl gene IDs. Haplosufficiency scores and ranks were then obtained for each ‘known’ gene. A consolidated dataset was then created (Data S1), containing Ensembl gene name, disease status, paralog status and both haplosufficiency score and rank, which provided the basis for the initial analysis of gene age, paralog status, disease status and haplosufficiency.

In order to investigate asymmetry the initial dataset was expanded. Four further datasets were compiled from data obtained the above sources, specifically pertaining to genes in the differing types of duplication pairs, including genes identified as [ohnolog]SSDs as [ohnolog-type] (Data S3:S5), and the other pertaining to genes in terminal paralog pairs (Data S2). For these files comparisons were made on a per pair basis. Each gene in the pair was named Conserved ‘C’ or Diverged ‘D’, where gene C had the shortest distance to root, and was therefore considered the most conserved, and gene D the greatest distance, and therefore considered the most diverged. Disease association per pair was then calculated, and disease associations previously listed as ‘both’ were considered dominant.

To validate the asymmetry between the pairs functional dissimilarity was calculated and added to the data. In order to do this the complete list of human GO terms was downloaded from the gene ontology consortium ((Ashburner et al., 2000), 31/10/2018), these were then stored for each gene, and compared between genes in each pair. For each term not present in the other gene the functional divergence score was increased by one. Any pairs with a score of less than one was considered not to exhibit true asymmetry, and therefore dropped from the asymmetry data.

### Analysis of Disease status, paralog status, and haplosufficiency

All further analysis and image generation was achieved using R version 3.3.2 (2016-10-31) "Sincere Pumpkin Patch" (R Core Team, 2016), and the following libraries; ggplot2 (Wickham, H, 2009) ggfortify (Yuan T, Masaaki H, Wenxuan L, 2016); dplyr (Wickham et al., 2017); lattice (Sarkar, 2017); plyr (Wickham, 2016); raster (Hijmans et al., 2017); gridExtra (Auguie and Antonov, 2017); tidyr (Wickham et al., 2018a); cluster (Maechler et al., 2018); FactoMineR (Husson et al., 2018); Devtools (Wickham et al., 2018b); factoextra (Kassambara and Mundt, 2017), and, corrplot (Wei et al., 2017). R Scripts supplied for figures 1–5 as supplementary data (S6).

## Acknowledgments

AM-G and MR are funded by PhD studentships from the MRC (#1622139)) and BBSRC ((#BB/M011208/1) respectively. DLR is partially funded by the MRC (MC UU 1201412). MT thanks the Newlife Foundation (#14-15/15) for funding.

## Author contributions

AM-G conducted the research, co-wrote, reviewed and edited the manuscript, produced all figures in main text, and performed the statistical analyses.

MR developed the FFS gene dating method, collaborated in the investigation of disease-association and haploinsufficiency, and reviewed and edited the manuscript.

BK conducted initial research investigating gene age, haploinsufficiency and disease.

MT contributed knowledge of human genetic disease, and reviewed and edited the manuscript.

DLR conceived the project, supervised all aspects of the research, co-wrote, reviewed and edited the manuscript, and collaborated in the generation of all figures

## Supplementary Figures

**Fig. S1.**
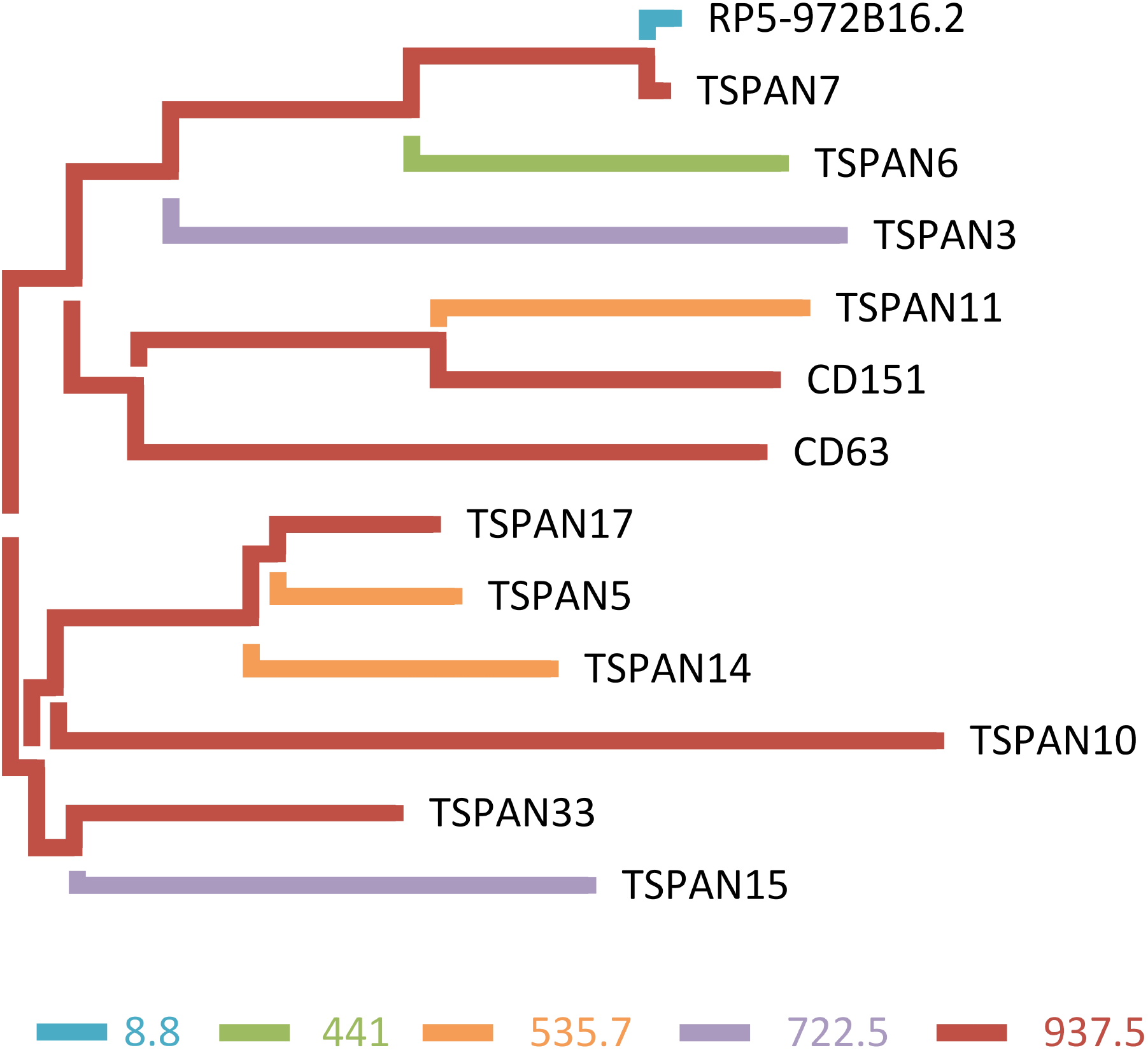
Allocation of duplication ages to an evolutionary tree of an example gene family, TSPAN, with gene age/ taxonomic levels indicated, see key, on the branch lengths.

**Fig. S2.**
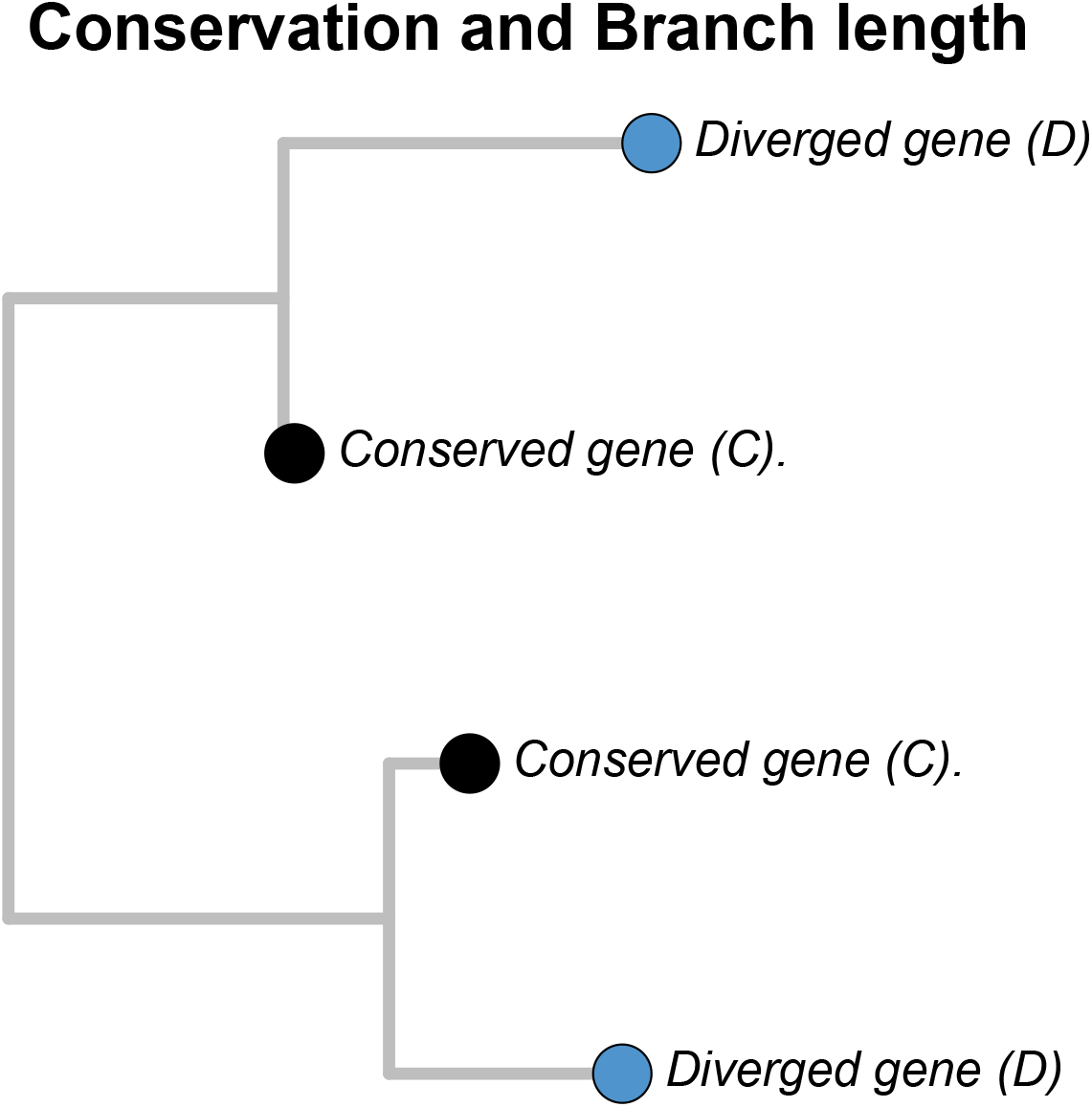
Example divergence of two gene pairs within a phylogenetic tree. Blue nodes show the more diverged genes in each pair, with the correspondingly longer branch length, and black nodes show the more conserved genes in each pair, with correspondingly shorter branch lengths.

**Table S1.** Absolute counts of genes with differing disease-associations and duplication histories within each FFS time-point.

| Duplicates |  |  |  |  |  |  | Singletons |  |  |  |  |  |
| --- | --- | --- | --- | --- | --- | --- | --- | --- | --- | --- | --- | --- |
| FFS | Duplicates | Dominant D | Recessive D | 'Both' D | Disease Unknown D | Undiseased D | Singltons | Dominant S | Recessive S | 'Both' S | Disease Unknown S | Undiseased S |
| 0 Homo Sapiens | 625 | 9 | 12 | 3 | 147 | 454 | 625 | 9 | 12 | 3 | 147 | 454 |
| 8.8 Homininae | 152 | 2 | 1 | NA | 49 | 100 | 152 | 2 | 1 | NA | 49 | 100 |
| 15.7 Hominidae | 58 | 1 | NA | NA | 20 | 37 | 58 | 1 | NA | NA | 20 | 37 |
| 20.4 Hominoidea | 56 | 2 | 1 | NA | 26 | 27 | 56 | 2 | 1 | NA | 26 | 27 |
| 29 Catarrhini | 242 | 2 | 3 | 3 | 99 | 135 | 242 | 2 | 3 | 3 | 99 | 135 |
| 42.6 Simiiformes | 158 | 1 | 3 | 1 | 67 | 86 | 158 | 1 | 3 | 1 | 67 | 86 |
| 65.2 Haplorrhini | 13 | NA | 1 | NA | 6 | 6 | 13 | NA | 1 | NA | 6 | 6 |
| 74 Primates | 33 | NA | 1 | NA | 11 | 21 | 33 | NA | 1 | NA | 11 | 21 |
| 92.3 Euarchontoglires | 46 | NA | 2 | NA | 13 | 31 | 46 | NA | 2 | NA | 13 | 31 |
| 100 Boreoeutheria | 82 | NA | NA | NA | 26 | 56 | 82 | NA | NA | NA | 26 | 56 |
| 104.2 Eutheria | 1868 | 11 | 8 | 3 | 278 | 1568 | 1868 | 11 | 8 | 3 | 278 | 1568 |
| 162.6 Theria | 473 | 6 | 4 | 1 | 122 | 340 | 473 | 6 | 4 | 1 | 122 | 340 |
| 167.4 Mammalia | 351 | 13 | 12 | 6 | 132 | 188 | 351 | 13 | 12 | 6 | 132 | 188 |
| 296 Amniota | 394 | 11 | 9 | 6 | 104 | 264 | 394 | 11 | 9 | 6 | 104 | 264 |
| 371.2 Tetrapoda | 110 | 6 | 7 | 1 | 57 | 39 | 110 | 6 | 7 | 1 | 57 | 39 |
| 414.9 Sarcopterygii | 143 | 5 | 4 | NA | 40 | 94 | 143 | 5 | 4 | NA | 40 | 94 |
| 441 Eutelestomi | 2743 | 185 | 200 | 64 | 1626 | 668 | 2743 | 185 | 200 | 64 | 1626 | 668 |
| 535.7 Vertebrata | 2200 | 160 | 211 | 67 | 1336 | 426 | 2200 | 160 | 211 | 67 | 1336 | 426 |
| 722.5 Chordata | 917 | 55 | 89 | 32 | 521 | 220 | 917 | 55 | 89 | 32 | 521 | 220 |
| 937.5 Bilateria | 2896 | 262 | 331 | 79 | 1745 | 479 | 2896 | 262 | 331 | 79 | 1745 | 479 |
| 1215.8 Opisthokonta | 769 | 64 | 104 | 18 | 493 | 90 | 769 | 64 | 104 | 18 | 493 | 90 |

**Table S2.**
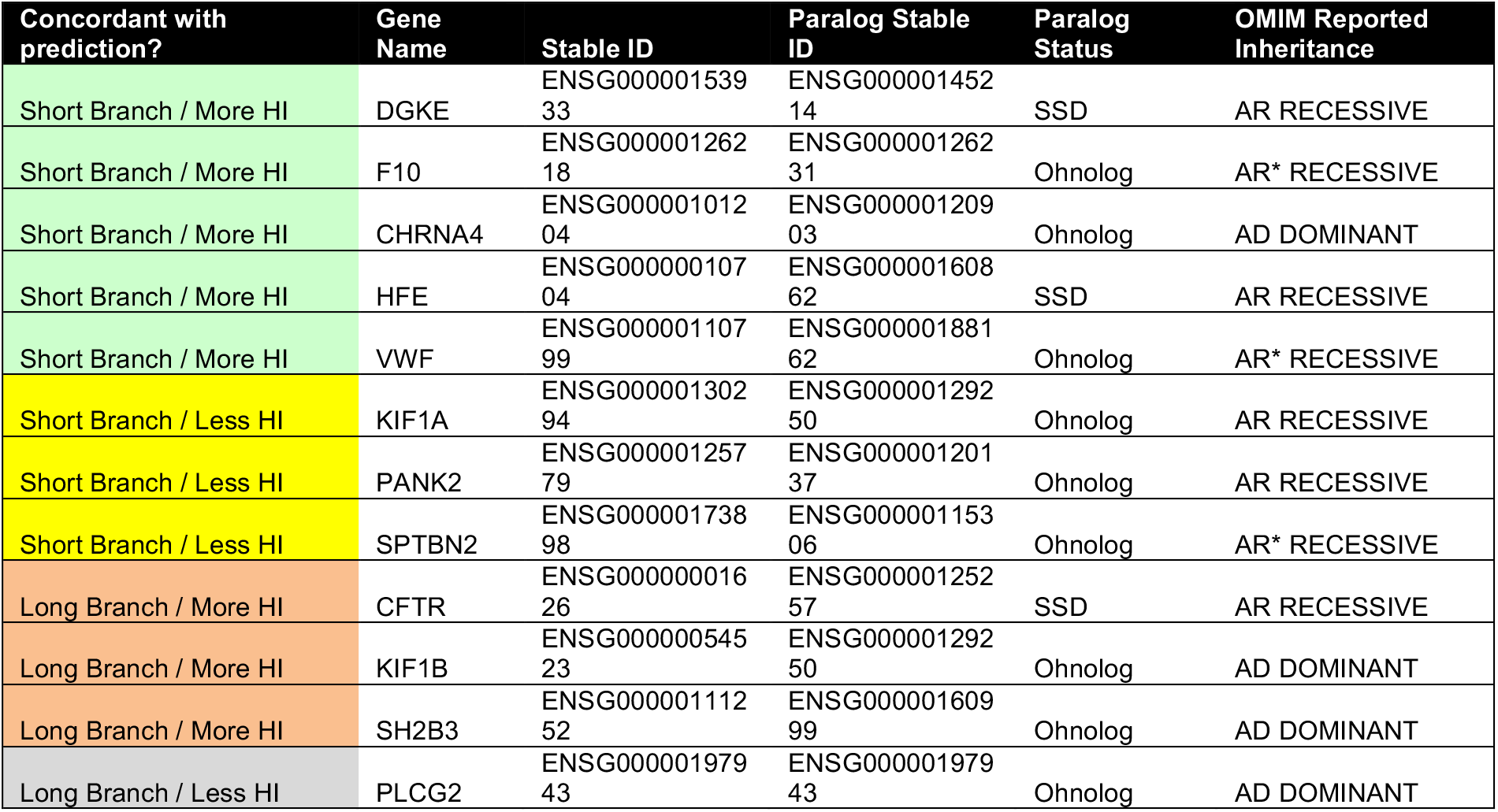
Summary of conservation and haploinsufficiency asymmetry in genes obtained from Bastarache *et al* (Bastarache et al., 2018). Column 1 indicates concordance with the predictions proposed within this manuscript.

